# Induction of a SALL4-dependency for targeted cancer therapy

**DOI:** 10.1101/2020.07.10.197434

**Authors:** Junyu Yang, Chong Gao, Miao Liu, Zhiyuan Chen, Yao-Chung Liu, Junsu Kwon, Jun Qi, Xi Tian, Alicia Stein, Yanjing Liu, Nikki R. Kong, Yue Wu, Shenyi Yin, Jianzhong Jeff Xi, Hongbo Luo, Leslie E. Silberstein, Julie A. I. Thoms, Ashwin Unnikrishnan, John E. Pimanda, Daniel G. Tenen, Li Chai

## Abstract

Oncofetal protein SALL4 is critical for tumor cell survival, making it a promising target in cancer therapy. However, it is detectable only in a subset of cancer patients, which limits the therapeutic impact of a SALL4 targeted therapy. Here we report that SALL4 can be activated and/or upregulated pharmacologically by hypomethylating agents, such as 5-Aza-2’-deoxycytidine (DAC), which are used clinically, and that SALL4 negative cancer cells become SALL4 dependent following exogenous expression of SALL4. In addition, the histone deacetylase inhibitor Entinostat (ENT) negatively regulates SALL4 expression by upregulating miR-205. Both ENT and miR-205 treatment induced cell apoptosis, rescuable by SALL4 expression or miR-205 inhibition. Finally, DAC pre-treatment sensitizes SALL4 negative cancer cell lines to ENT both in culture and *in vivo* by upregulating SALL4. Overall, we propose a framework whereby the scope of targeted therapy can be expanded by sensitizing cancer cells to treatment by target induction and engineered dependency.

**Significance:** This proof of concept study demonstrates that targeted cancer therapy can be achieved by inducing a targetable gene establishing a survival-dependency for cancer cells. For SALL4, sequential treatment of DAC and ENT could expand the scope of SALL4 targeted cancer therapy.

## Introduction

SALL4 is a zinc finger transcription factor that belongs to the spalt-like (SALL) gene family. SALL4 plays a significant role in the maintenance of self-renewal and pluripotency of embryonic stem cells(*1, 2*). While SALL4 expression is silenced in most healthy adult tissues, it is re-expressed in various cancers and is regarded as an important biomarker for poor outcomes (*3-6*). Targeting SALL4 by RNA interference or peptides has been shown to be effective against a wide range of cancers, including lung cancer(*7*), endometrial cancer(*8*), gastric cancer(*9*), and liver cancer(*10*). However, the fact that SALL4 is only positive in a sub-set of cancer patients limits the application of SALL4-based cancer treatment(*11-13*), which is broadly true for all targeted therapies.

The function and downstream targets of SALL4 have been explored and identified(*9, 14, 15*). However, the regulation of SALL4 remains unexplored. Methylation of the SALL4 locus has been reported to be negatively associated with SALL4 expression (*16*). Hypomethylation agents (HMA), such as 5-aza-2’-deoxycytidine (DAC) and 5-azacytidine (5-Aza), are FDA-approved agents used to treat myelodysplastic syndrome (MDS) and acute myeloid leukemia (AML)(*17-19*), functioning at least in part through inhibition of DNA methyltransferases (DNMTs)(*20*). The effects of HMA on SALL4 expression have not been carefully examined. SALL4 functions as a gene repressor by interacting with the Nucleosome Remodeling Deacetylase (NuRD) complex, in which histone deacetylases (HDACs) are critical components(*21*). We have previously shown that the gene signatures of the HDAC inhibitor Entinostat (ENT) and SALL4 were correlated using the connectivity map tool(*7*). We rationalized at that time that by blocking HDAC enzymatic activity, ENT functions as a SALL4 inhibitor by a mechanism akin to a SALL4 peptide blocker(*7*). Intriguingly, we also noticed that SALL4 protein level was significantly decreased upon ENT treatment, while the mechanism remained unclear. This prompted us to examine whether ENT can potentially function as a SALL4 inhibitor drug by modulating its expression possibly by up regulating a regulatory RNA.

MicroRNAs (miRNAs) are a class of small non-coding RNAs that act as post-transcriptional regulators by inducing mRNA degradation or translation repression of their targets through directly binding to the 3’untranslated region (3’UTR)(*22-24*). MiRNAs have been demonstrated to participate in various biological processes, such as self-renewal, cell proliferation, cell cycle, migration and apoptosis(*25-27*). Dysregulation of miRNAs are frequently observed during the development and tumorigenesis(*28-30*). Several mechanisms have been identified contributing to the aberrant expression of miRNAs in cancers, in which epigenetic changes exhibit a significant role(*31-33*).

In this study, we demonstrate that SALL4 expression was found to be negatively correlated with its DNA methylation and could be upregulated by DAC treatment. After exogenous expression, SALL4-negative cancer cells could become SALL4-dependent. Increased SALL4 expression can enhance cancer cells sensitivity to ENT, a SALL4 inhibitor that regulates SALL4 expression post transcriptionally through miRNA-205. We have further evaluated the potential of DAC plus ENT based therapy to expand SALL4-targeted therapy in SALL4 negative cancers using cell culture and *in vivo* xenotransplantation models.

## Results

### A SALL4-dependency can be established in SALL4 negative cancer cells

Previously, we and others have shown that SALL4 is critical for cancer cell survival in SALL4 positive tumors(*7, 8, 10*). Using the Dependency Map (DepMap) analytical tool, we observed that cancer cells with high SALL4 expression exhibited more dependency on SALL4 compared to those with low SALL4 expression (Supplement Figure 1A). However, it remains unknown whether SALL4-dependency can be established in SALL4 negative cancers. To answer this question, SALL4 was stably introduced into SALL4 negative H1299 lung cancer cells by retroviral transduction (designated H1299-SALL4). Intriguingly, treatment with shRNA against SALL4 led to significant inhibition of cell viability and induction of apoptosis in H1299-SALL4, but not in control H1299-GFP cells (Figure 1A-D). Loss of viability and induction of apoptosis in H1299-SALL4 following SALL4 repression mirrored the phenotype observed when knocking down SALL4 in H661 lung cancer cells, which express endogenous SALL4 (Figure 1E-H). This suggests that SALL4 negative cancer cells can become addicted to SALL4 once expressed. A similar SALL4-dependency was also observed in the SALL4 positive liver cancer cell line SNU398 (Supplement Figure 1B) and paired isogenic SALL4 negative liver cancer SNU387 cells with and without SALL4 expression (Supplement Figure 1C).

**Figure 1.**
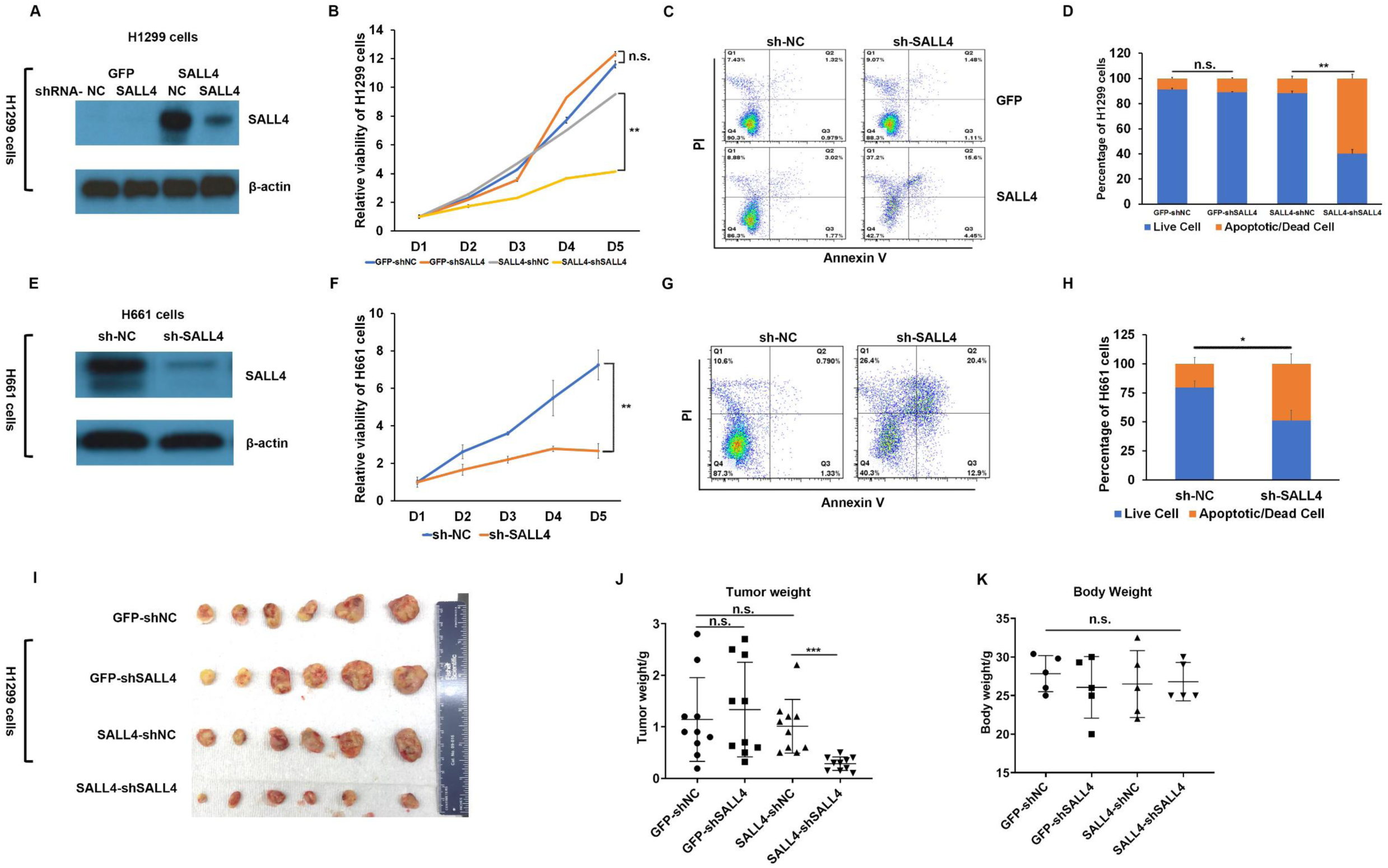
Engineered SALL4-dependency in SALL4-negative cancer cells. (A) Western blot of SALL4 protein in GFP control or exogenous SALL4-expressing H1299 cells 48 hours after transfection of negative control or SALL4 shRNA, β-actin was used as normalized control. (B) Cell viability of GFP control or SALL4-expressing H1299 cells after the indicated treatments. (C) Flow cytometry plots showing Annexin/PI staining of GFP control or SALL4-expressing H1299 cells after the indicated treatments. (D) Percent apoptotic (Annexin^+^) and dead cells (Annexin^-^ /PI^+^) from (C). (E) Western blot of SALL4 protein in H661 cells 48 hours after transfection of negative control or SALL4 shRNA. β-actin was used as normalized control. (F) Cell viability of H661 cells after indicated treatment. (G) Flow cytometry plots showing Annexin/PI following the indicated treatment. (H) Percent apoptotic and dead cells from (G). (I) Representative images of tumors harvested from mice 30 days following sub-cutaneous injection of H1299 cells expressing the indicated vectors. (J) Tumor weight and (K) body weight from indicated groups (N=5). n.s. means P>0.05, *P<0.05, **P<0.01, ***P<0.001, N=3.

To explore a potential mechanism for SALL4-mediated dependency in cancer cells, RNA-sequencing was performed in SNU387 cells with or without SALL4 overexpression. Among 1857 genes with significant up or down regulation (over 2-fold) after SALL4 was introduced (SNU387-SALL4 vs SNU387-EV), there were 613 genes overlapping with SALL4 positive SNU398 cells (Supplement Figure 1D and E), indicating an important role of SALL4 in cancer cell reprogramming. GO analysis indicated that these genes were enriched in biological processes such as nuclear DNA replication, cell cycle DNA replication, and different types of cell morphogenesis and development (Supplement Figure 1F).

In addition, inhibition of SALL4 in a xenograft mouse model of H1299-SALL4 also led to a highly significant reduction of tumor growth (Figure 1I-K). Taken together, these results demonstrated that expression of SALL4 in negative cancer cells sensitizes these cells to SALL4 targeting with inhibition of cell growth both in cell culture and *in vivo*.

### Upregulation of SALL4 post HMA treatment

We next evaluated whether inducing SALL4 pharmacologically create a dependency that could be therapeutically targeted. SALL4 expression is negatively correlated with DNA methylation at exon 1 (manuscripts submitted and (*16*). We therefore explored the relationship between SALL4 RNA expression levels and density of DNA methylation at exon 1 in 386 cancer cell lines (CCLE database) and in 722 primary tumors (TCGA database, including lung adenocarcinoma and squamous cell carcinoma, colon adenocarcinoma, hepatocellular carcinoma, and gastric adenocarcinoma). As shown in Figure 2A and B, compared to the high methylation (>75% methylation) groups, the expression of SALL4 was significantly increased (4-fold in cell lines, and 2.4-fold in primary cancer patients) in the low methylation (<25% methylation) groups. 5-azacitidine has been approved by the FDA for the treatment of myelodysplasia, chronic myelomonocytic leukemia (CMML) and acute myeloid leukemia. Following six cycles of 5-azacitidine therapy, SALL4 expression was increased in bone marrow mononuclear cells in 7/13 patients, compared with pre-treatment levels (Figure 2C).

**Figure 2.**
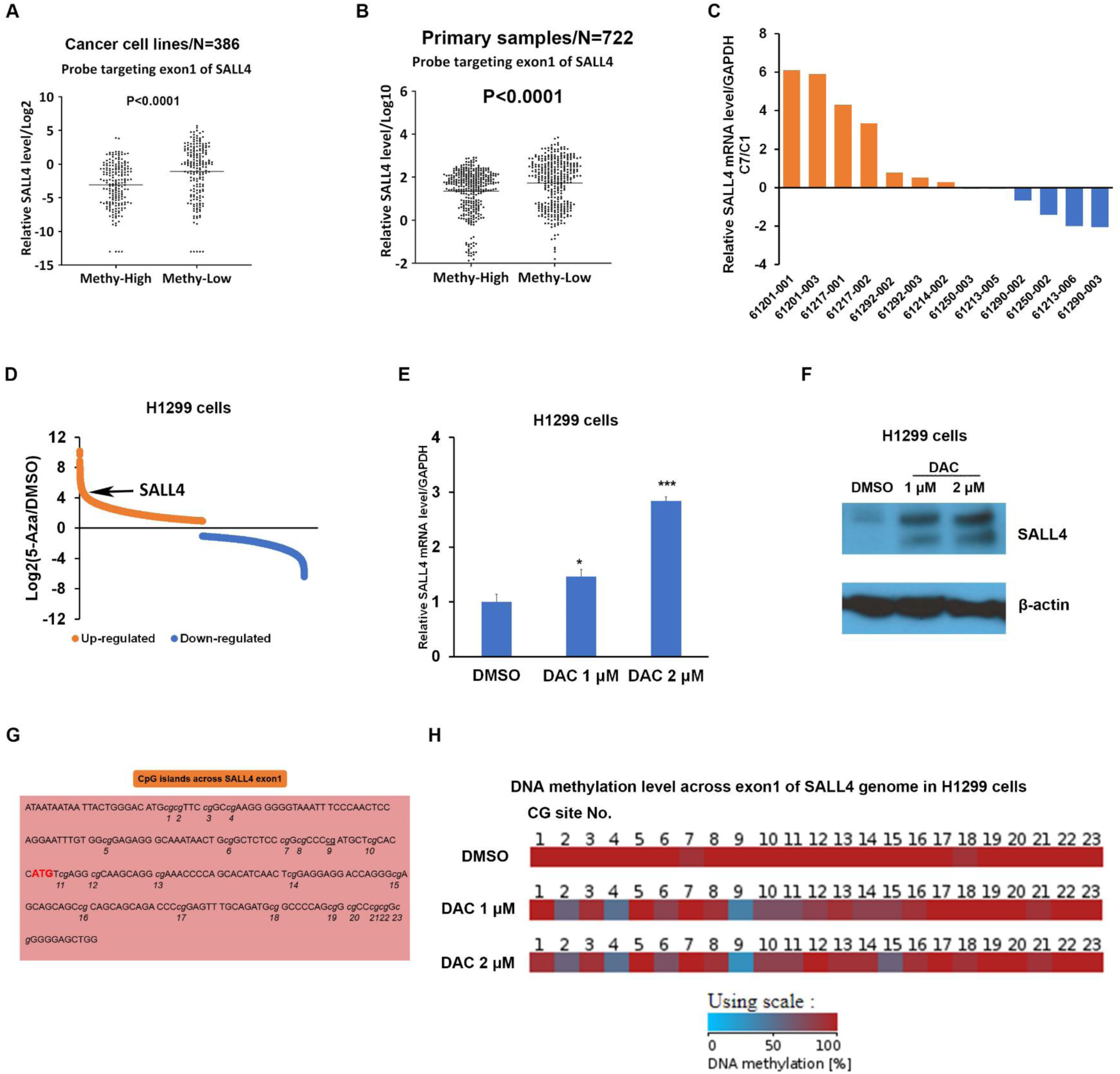
Upregulation of SALL4 by hypomethylating agents. (A) SALL4 expression levels in cancer cell lines with either high-(>75%) and low-(<25%) CpG methylation at exon 1, data generated from CCLE database. (B) SALL4 expression levels in primary tumors from lung adenocarcinoma and squamous cell carcinoma, colon adenocarcinoma, hepatocellular carcinoma, and gastric adenocarcinoma with either high-(>75%) and low-(<25%) CpG methylation at exon 1, data generated from TCGA database. (C) Fold change of SALL4 expression measured by real-time PCR from patients’ bone marrow mononuclear cells after six cycles of 5-Aza treatment (C7D1/C1D1), GAPDH was used for a normalization control. (D) RNA-seq data of H1299 cells after treatment of 1 μM 5-Aza or DMSO, data generated from GSE5816. (E) mRNA and (F) protein levels of SALL4 in H1299 cells after 5 days treatment of DMSO or DAC. (G) Map of CpG islands tested within the first exon of SALL4 locus. (H) Methylation levels of CpG islands within the first exon of SALL4 genome in H1299 cells after indicated treatment. *P<0.05, ***P<0.001, N=3.

Upon data mining from a published gene expression database (*34*), we noticed that DAC treatment led to significantly increased SALL4 RNA expression levels in H1299 cells (Figure 2D, generated from GSE5816). We then confirmed that SALL4 mRNA and protein expression in H1299 cells was indeed significantly up-regulated upon DAC treatment (Figure 2E and F) along with decreased methylation level at exon 1 (Figure 2G and H). Similar results were also observed in the SALL4 negative liver cancer cell line SNU387 and leukemia cell lines K562 and HL60 (manuscripts submitted). This contrasts with the results of H661 cells (high in endogenous SALL4) after DAC treatment, in which no change in SALL4 mRNA (Supplement Figure 2A) or protein (Supplement Figure 2B) expression was observed. Together, these results indicated that HMA treatment can upregulate SALL4 in negative cancer cells.

### Entinostat (ENT) represses SALL4 expression post transcriptionally

We next reviewed existing drugs that could target SALL4 in cancer. We previously identified Entinostat (ENT) as a SALL4 inhibitor, and its treatment triggered decreased SALL4 protein expression through an unknown mechanism(*7*). To elucidate the effects of ENT on SALL4, we first examined SALL4 protein expression levels at various time points post treatment. SALL4 protein expression in SALL4+ H661 cells was significantly decreased within 48 hours of ENT treatment at a concentration of 2.5 μM (Figure 3A), along with significantly decreased mRNA expression (Figure 3B and Supplement Figure 3A). Surprisingly, we noticed that SALL4 pre-mRNA level was increased after ENT treatment, in contrast to its mature mRNA (Figure 3C). Similar results were observed in SALL4+ SNU398 cells after ENT treatment (Supplement Figure 3B and C). In addition, we also examined the methylation status at exon-1 before and after DAC treatment. As shown in Supplement Figure 3D, compared to control DMSO treatment, there was no significant change in the average methylation levels after ENT treatment at different time points. Meanwhile, a significant up-regulation was observed in H3K27 acetylation (H3K27ac), which in general correlates with open chromatin and gene activation, at the SALL4 promoter region (Supplement Figure 3E). These results indicate that ENT-induced SALL4 repression was probably at a post transcriptional level.

**Figure 3.**
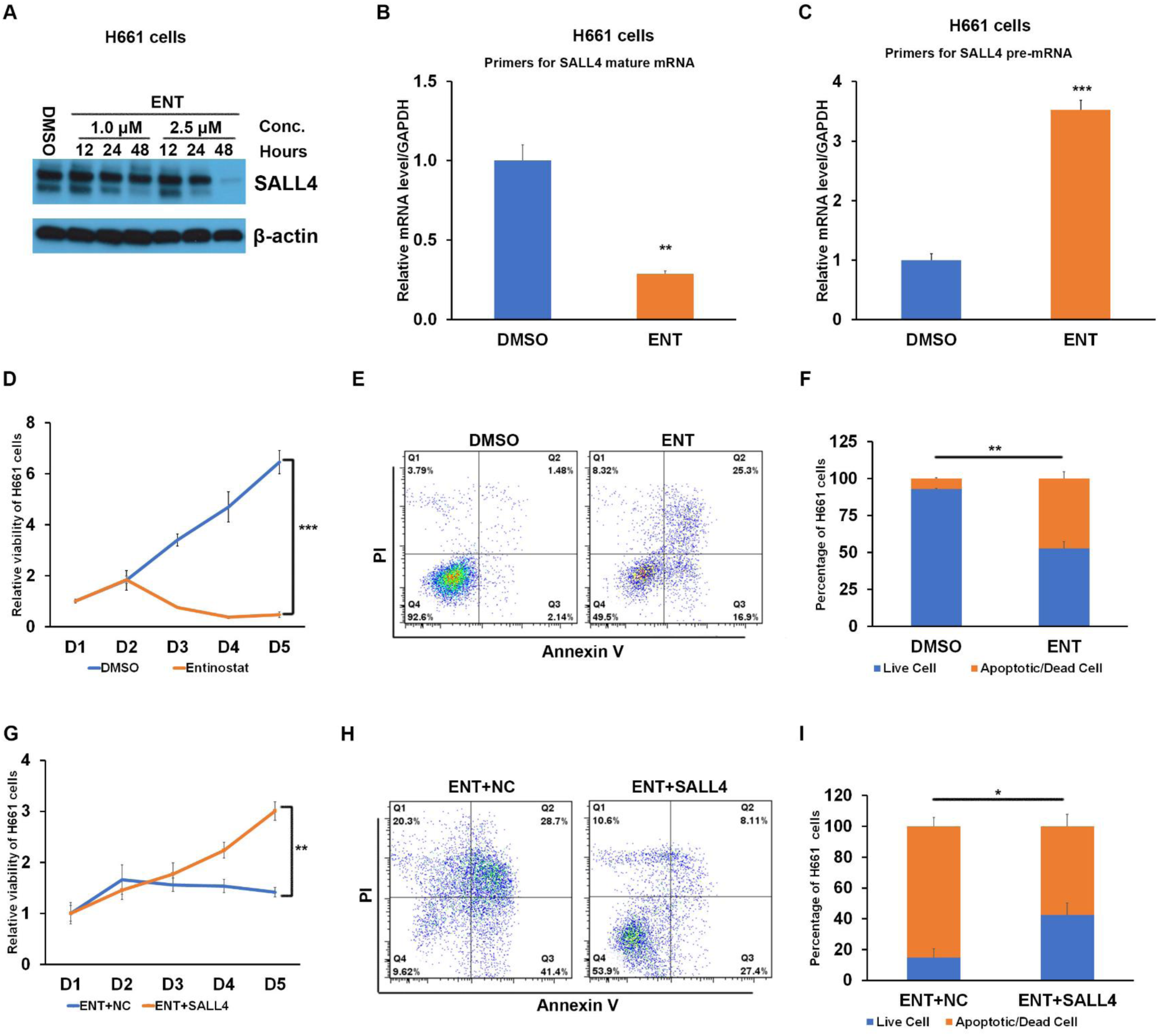
ENT treatment decreased cell viability and induced cell apoptosis via inhibition of SALL4. (A) Protein and (B) mRNA levels of SALL4 in H661 cells after the indicated treatment of ENT or DMSO control. (C) Pre-mRNA level of SALL4 in H661 cells after 48 hours treatment of ENT or DMSO control. (D) Cell viability of H661 cells after indicated treatment for 5 days. (E) Flow cytometry of H661 cells after indicated treatment. (F) Percent apoptotic and dead cells in (E). (G) Cell viability of ENT pre-treated H661 cells after transfection of negative control or SALL4 overexpression vector. (H) Flow cytometry of ENT pre-treated H661 cells after indicated treatment. (I) Percent apoptotic and dead cells in (H). *P<0.05, **P<0.01, ***P<0.001, N=3.

Furthermore, the IC_50_ of lung cancer cell lines for ENT was negatively correlated with their endogenous SALL4 levels (Supplemental Figure 3F). ENT treatment of H661 cells led to significantly decreased cell growth (Figure 3D) and increased apoptosis (Figure 3E and F). To confirm the ENT-mediated cellular effects were through SALL4 inhibition, we overexpressed SALL4 in H661 cells before ENT treatment. As shown in Figure 3G-I, SALL4 overexpression rescued the loss of viability and induction of apoptosis triggered by ENT treatment, compared to the control group. Taken together, our data suggested that ENT could suppress cell growth and induce apoptosis in cancer cells by post-transcriptional inhibition of SALL4.

### ENT-induced SALL4 inhibition is mediated by microRNA miR-205

As a class of post-transcriptional regulators, miRNAs were considered as the potential mediator in ENT-induced SALL4 inhibition. MiRNA-sequencing was performed using H661 cells after 8 hours of ENT treatment, at a time point much earlier than when SALL4 RNA expression changes were observed (Figure 4A). Using a 2-fold change as a threshold, 38 miRNAs were significantly up-regulated by ENT treatment, and 32 miRNAs were significantly decreased (Supplemental file1). After overlapping with the Targetscan database, miR-205 was identified as the top potential candidate that was significantly enhanced by ENT treatment and predicted to target SALL4 (Figure 4B). MiR-205 is highly conserved among species. In humans, it is located at the intronic region of the host gene on chromosome 1. Pathway analysis of predicted targets showed that miR-205 was potentially involved in tumor development, cell cycle, and DNA repair (Supplement Figure 4A). The expression level of miR-205 was confirmed to be significantly up-regulated following ENT treatment (Figure 4C). Moreover, as shown in Figure 4D, ENT significantly increased the H3K27ac level at the miR-205 promoter region, which correlated with increased expression of miR-205.

**Figure 4.**
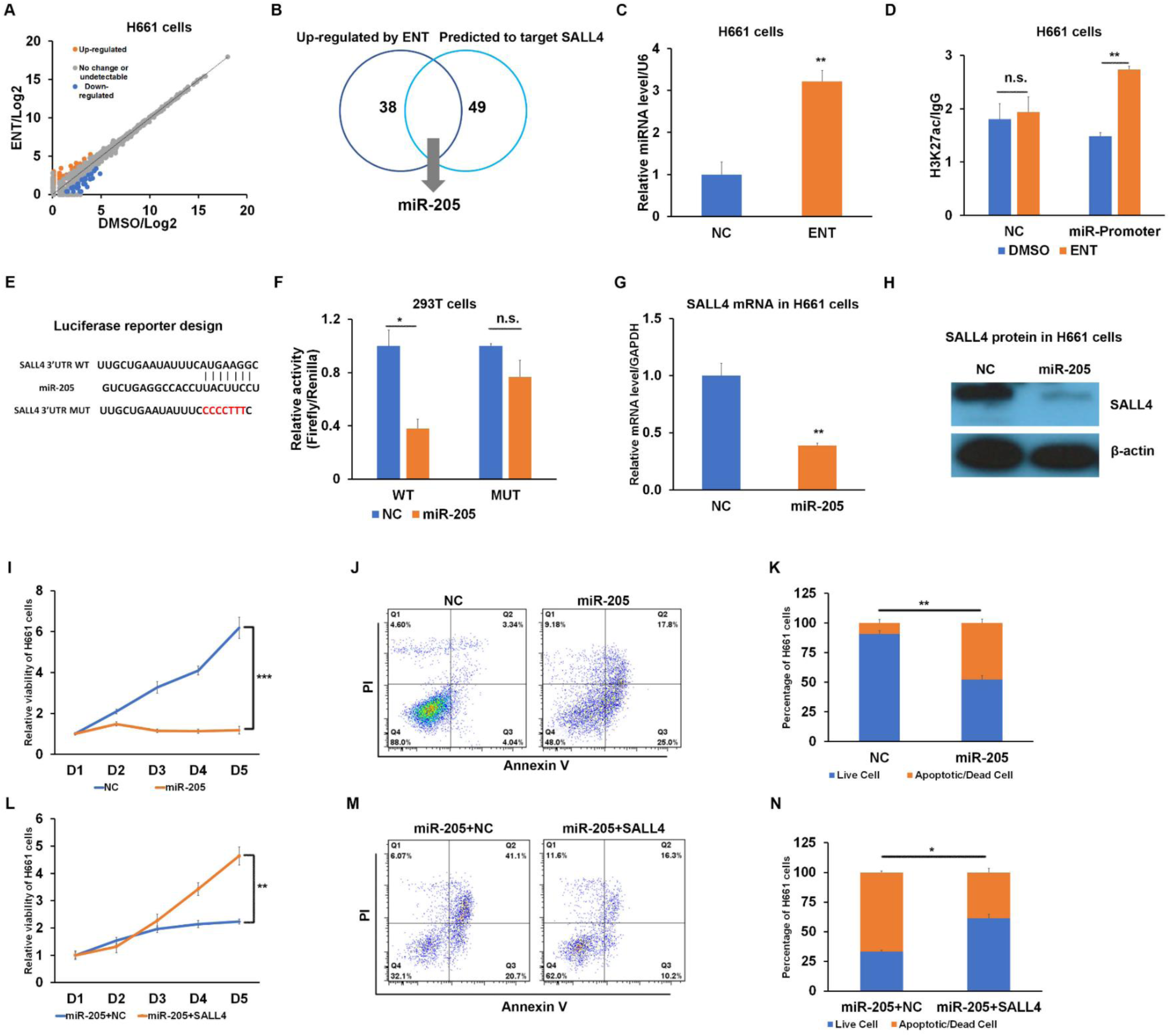
MiR-205 involved in ENT-induced SALL4 inhibition. (A) miRNA sequencing results of H661 cells after 8 hours treatment with 2.5 μM ENT or DMSO. (B) miR-205 was up-regulated by ENT and predicted to target SALL4 by the Targetscan database. (C) Real-time PCR of miR-205 expression in H661 cells after the indicated treatment for 8 hours. U6 was used as a normalization control. (D) ChIP-qPCR of the miR-205 promoter region and non-specific control region after ENT treatment. IgG was used as a normalization control. (E) Sequence of wild-type SALL4 3’UTR with miR-205 binding site or mutant one without binding site. (F) Luciferase reporter result in 293T cells after transfection with the indicated vectors. (G) mRNA and (H) protein levels of SALL4 in H661 cells after transfection with miR-205 or non-specific control. (I) Cell viability of H661 cells after indicated treatment for 5 days. (J) Flow cytometry of H661 cells after the indicated treatment. (K) Percent apoptotic and dead cells in (J). (L) Cell viability of ENT pre-treated H661 cells after transfection of miR-205 or non-specific control. (M) Flow cytometry of ENT pre-treated H661 cells after indicated treatment. (N) Percent apoptotic and dead cells in (M). n.s. means P>0.05, *P<0.05, **P<0.01, ***P<0.001, N=3.

Next, we examined whether miR-205 could target SALL4 to repress its expression. As shown in Figure 4E and F, luciferase reporter assays demonstrated that miR-205 could directly target SALL4 wild-type mRNA and inhibit luciferase activity, while it had no significant effect on mutant constructs lacking a binding site for miR-205. Meanwhile, overexpression of miR-205 in H661 cells significantly repressed SALL4 expression at both mRNA and protein levels compared to control (Figure 4G and H). Immunofluorescent staining of H661 cells after transfection with GFP-tagged scramble or miR-205 plasmid also showed that SALL4 level was significantly lower in cells with miR-205 overexpression (Supplement Figure 4B). Furthermore, similar to ENT treatment, miR-205 overexpression in H661 cells also repressed cell number (Figure 4I) and induced apoptosis (Figure 4J and K), which was rescued by SALL4 overexpression (Figure 4L-N). Inhibition of miR-205 by its antisense oligo (miR-205AS) was sufficient to restore SALL4 expression in the presence of ENT treatment (Supplement Figure 4C and D) and prevented cells from ENT-induced growth repression and apoptosis (Supplement Figure 4E-G). Taken together, these data support the premise that miR-205 was responsible for ENT-mediated SALL4 inhibition.

### DAC pre-treatment can prime SALL4 negative cells to be targetable by the SALL4 inhibitor ENT

Based on the results shown above that SALL4 could be induced by DAC and targeted by ENT, we further tested whether DAC-primed SALL4 negative cancer cells could become sensitive to ENT. Five days of DAC treatment at a concentration of 1 or 2 μM was performed in both H661 and H1299 cells followed by the evaluation of drug sensitivity to ENT (Figure 5A). As shown in Figure 5B, there was no significant change of the IC_50_ of ENT for SALL4 positive H661 cells. In contrast, upregulation of SALL4 in originally negative H1299 cells rendered them to be more sensitive to ENT, with a significantly decrease of IC_50_ from 11.77 μM to 1.42 μM (Figure 5C), similar to the IC_50_ level of H661. Notably, ENT treatment also induced miR-205 expression in DAC-treated H1299 cells (Supplement Figure 5A), suggesting the same mechanism for SALL4 targeting. Moreover, similar results were also observed in SALL4 negative liver cancer SNU387 cell line (Supplement Figure 5B), whereby pretreatment of SNU387 can sensitize these cells to ENT with reduced IC_50_, and leukemia K562 and HL-60 cell lines, whereby DAC priming followed by ENT treatment in these cells led to significantly decreased cell viability (Supplement Figure 5C and D).

**Figure 5.**
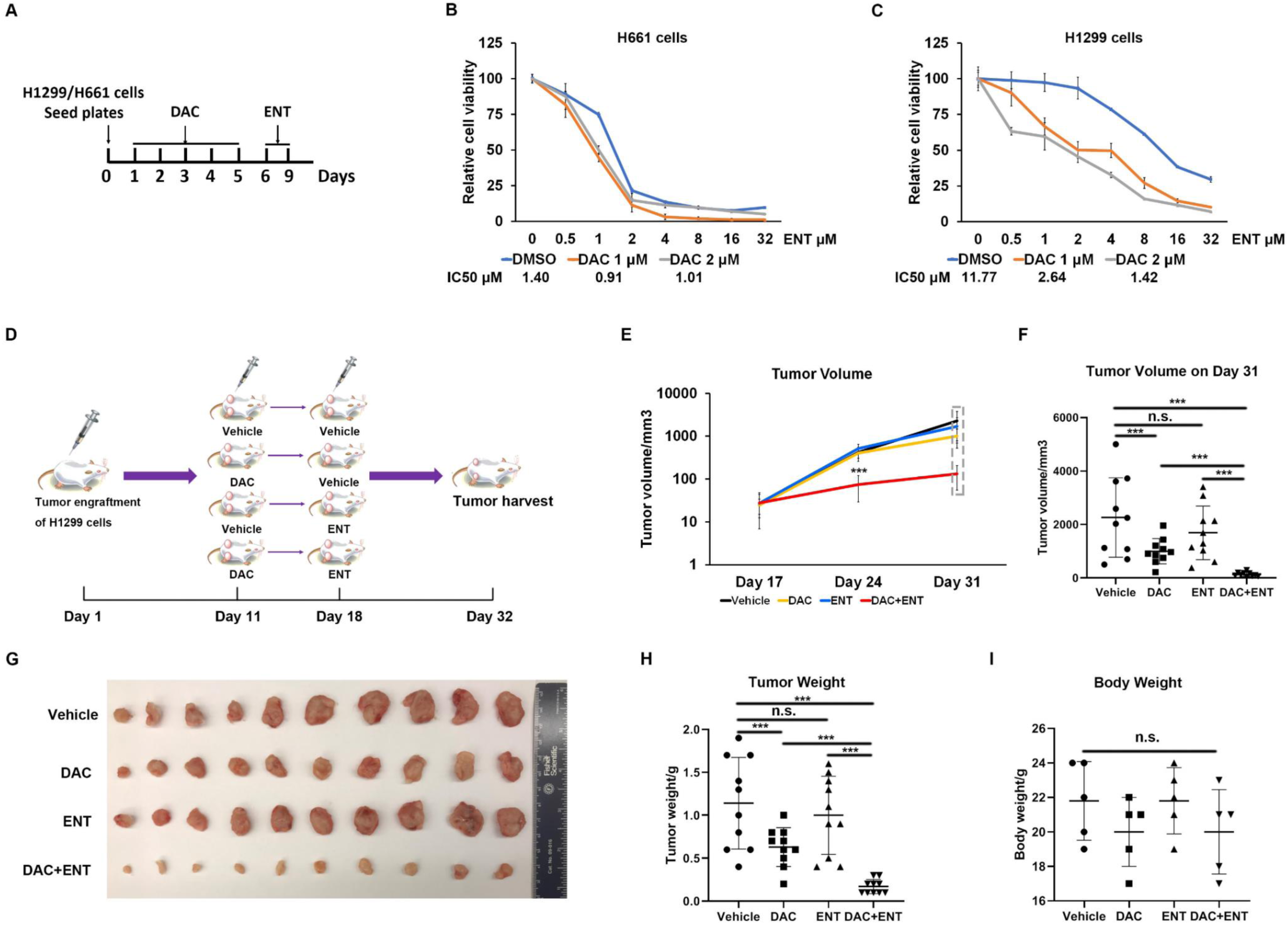
DAC/ENT combination treatment targeted SALL4 negative cancer cells. (A) Schema of DAC and ENT combination treatment of H1299 or H661 cells. (B) Relative cell viability and IC50 of DMSO or DAC pre-treated H661 cells following ENT therapy. (C) Relative cell viability and IC50 of DMSO or DAC pre-treated H1299 cells following ENT therapy. (D) Schema of sequential drug treatment in a murine xenograft model. (E) Tumor growth measured at different time points. (F) Tumor volume measured at the end point from the indicated treatment groups. (G) Image of tumors harvested from mice with H1299 cells xenotransplantation after indicated treatment. (H) Tumor weights and (I) body weights from indicated groups. **P<0.01, ***P<0.001, N=3.

We next evaluated whether this sequential drug treatment strategy would work *in vivo*. H1299 cells were implanted into non-obese diabetic scid gamma (NSG) mice and treated with sequential injections of DAC and ENT both at a dose of 2.5 mg/kg according to published protocols (Figure 5D)(*35-37*). As shown in Figure 5D-E, tumors were monitored and measured on day 17, 24, and day 31, and were harvested on day 32 after various treatments. Following DAC and the first 5 doses of ENT injection at day 24, the sizes of tumors from mice received both DAC and ENT treatment were significantly smaller than other groups (Figure 5E). At the end point of the study, the size and weight of tumors from mice that received DAC+ENT treatments were remarkably smaller than vehicle-, DAC-, and ENT-treated controls (Figure 5F-H). The body weights of all mice after different treatments exhibited no change (Figure 5I). Collectively, the results from cell culture and xenograft tumor models demonstrate show that DAC treatment can prime SALL4 negative cancer cells to be targeted by ENT, a SALL4 inhibitor.

## Discussion

Targeted cancer therapy has been of interest and a focus of research and development since the discovery of the first oncogenes in the 1960s(*38*), followed by the identification of tumor suppressor genes, mutated growth factor receptors, overexpressed signaling pathways, and mutated transcription factors. Targeted therapy has resulted in major advances in both solid and hematopoietic malignancies, but its application is limited to the fraction of patients with tumors that are positive for the target. As an important oncofetal gene, SALL4 is silenced in most healthy adult tissues but is reactivated in some tumors and contributes significantly to the survival of tumor cells. These features of SALL4 make it a promising target for cancer treatment. Studies have utilized RNA interference methods or peptides against SALL4 to effectively inhibit tumor growth in cell culture and xenotransplant models (*8-10*), and drugs that can be used to target SALL4 specifically in cancer patients are being developed. However, as in all targeted therapies, SALL4-centered treatment is limited to patients with tumors that are SALL4 positive, which represent only a sub-set of all cancer patients (*11-13*).

We have previously observed that exogenous expression of SALL4 in negative liver cancer cells could enhance their sensitivity to oxidative phosphorylation inhibitors due to a shift in metabolism(*39*), suggesting SALL4 could reprogram cancer cells. In our current studies, we further observed that SALL4 negative cancer cells can become SALL4 dependent. Knocking down SALL4 by shRNA in lung cancer H1299-SALL4 or liver cancer SNU387-SALL4 cells with exogenous SALL4 expression significantly inhibited cell viability, and mirrored the phenotypes observed in paired H661 and SNU398 cancer cells with endogenous SALL4 expression. Xenotransplants also confirmed that by knocking down SALL4 in isogenic H1299-SALL4 cells led to decreased tumor growth *in vivo*.

In our previous studies, we have shown that the HDAC inhibitor ENT, functions as a SALL4 inhibitor based on a drug/gene signatures observed using the connectivity map as an analytical tool(*7*). In our current study, we have identified a mechanism by which ENT acts as a SALL4 inhibitor drug. We report that ENT negatively regulates SALL4 expression post transcriptionally by epigenetically up-regulating miR-205 expression. Importantly, the impact of ENT on cell viability and tumor growth was dependent on miR205 expression and SALL4 inhibition. A CpG region at the 5’UTR-Exon 1-Intron 1 of SALL4, when highly methylated, acts as negative regulatory region for SALL4 expression. We previously reported that HMA treatment could upregulate SALL4 in acute B cell lymphoblastic leukemia(*16*). Here we show that treatment with HMA results in hypomethylation of this region and increased SALL4 expression. Indeed, several SALL4 negative cancer cell lines display significantly increased sensitivity against ENT after DAC treatment. The degree of ENT sensitivity in these SALL4 negative cancer cells in which SALL4 re-expression had been pharmacologically engineered by prior HMA therapy was comparable to ENT sensitivity of SALL4 positive cancer cells.

This therapeutic effect of DAC plus ENT for SALL4 non-expressing and otherwise ENT insensitive tumors was confirmed in xenograft murine models. This observation supports a prior report that ENT in combination 5-AZA could repress tumor growth and ablate up to 75% of tumor mass in a murine lung cancer model(*40*). In clinical phase II studies, combination therapy with low dose 5-azacytidine and ENT were well tolerated and resulted in durable response or improved progression-free survival in patients with refractory advanced non-small cell lung cancer(*41*), advanced breast cancer(*42*), and recurrent metastatic colorectal cancer(*43*). It is possible that SALL4-mediated dependence plays a role in these studies, and we plan to investigate this in cancer patients receiving HMA and ENT in the future.

In conclusion, our study has demonstrated that induction/upregulation of SALL4 in SALL4 negative cancer cells can sensitize these cells to SALL4 targeting drug(s). More specifically, in this study we propose that hypomethylation treatment can pharmacologically reactivate/upregulate SALL4 in cancer cells and prime them to SALL4 inhibitor drug (ENT) treatment (Figure 6). Overall, our studies support a scenario in which tumors can be pharmacologically sensitized to targeted therapy when reactivating a target induces cancer cell addiction to this target.

**Figure 6.**
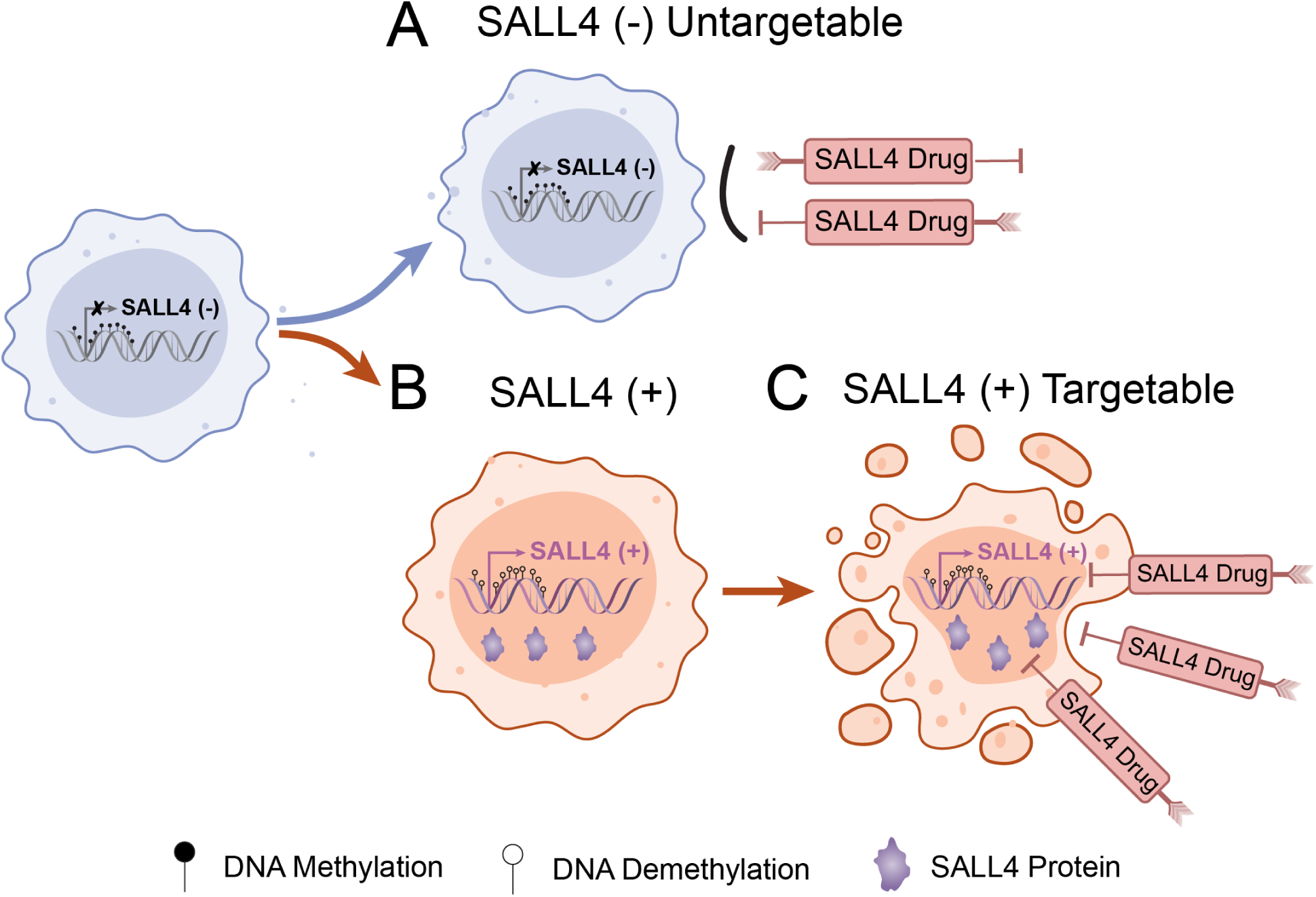
Induction of a SALL4-dependence for targeted cancer therapy: **(A)** SALL4 negative cells cannot be targeted by a drug that targets SALL4 specifically; **(B)** SALL4 negative cells can be induced to become SALL4 positive using genetic or pharmacological approaches; and then become targetable by SALL4 targeted therapies (C).

## Author contributions

J.Y. designed research, carried out experiments, analyzed data, prepared figures, and wrote the manuscript. C.G., Y.L., J.K., S.Y. JT, AU and JEP and A.S. carried out experiments and edited the manuscript. Z.C. performed the bioinformatics analysis on SALL4 expression and methylation. J.Q., N.R.K., Y.W., M.L., J.J.X., H.L. and L.E.S. reviewed the manuscript. D.G.T. and L.C. conceived of and supervised the project, designed experiments, and critically reviewed the manuscript.

### Acknowledgement

This study was supported by Singapore Ministry of Health’s National Medical Research Council (Singapore Translational Research (STaR) Investigator Award, D.G.T.; NMRC/OFIRG/0064/2017) and the Singapore Ministry of Education under its Research Centres of Excellence initiative; Singapore Ministry of Education Academic Research Fund Tier 3, grant number MOE2014-T3-1-006 (D.G.T); NIH/NCI Grant R35CA197697 and NIH/NCI CA66996 (D.G.T); as well as NIH/NHLBI Grant P01HL095489 and Xiu research fund (L.C.).

## Methods

### Cell culture and reagents

Lung cancer cell lines H661 and H1299, hepatocellular cancer cell lines SNU387 and SNU398, ovarian cancer cell line OV90, leukemic cell lines K562 and HL60 were cultured in RPMI 1640 medium (Thermo Fisher Scientific, Waltham, USA) supplied with 10% fetal bovine serum and 1% antibiotics (100 U/ml penicillin and 100 mg/ml streptomycin) at 37°C in 5% CO2. 293T cells were cultured in DMEM medium (Thermo Fisher Scientific, Waltham, USA) with the same condition. ENT was provided by Jun Qi’s lab from Dana Farber Cancer Institute. 5-aza-2’-deoxycytidine was purchased from Sigma-Aldrich (Cat. A3656, Louis, USA).

### Cell transfection

MiRNA negative controls, miR-205 mimics, and its inhibitor were purchased from Sigma-Aldrich (Louis, USA). shRNA against SALL4 was designed as previously described(*1*). MiRNAs or vectors were transfected in the amount of 2 μg for 6-well plate into indicated cells by Lipofectamine 2000 transfection reagent (Thermo Fisher Scientific, Waltham, USA) according to the manufacturer’s instructions. After 48 hours, the cells were used for subsequent experiments.

### Total RNA extraction and quantitative real-time PCR

Cells were dissolved in TRIzol reagent (Thermo Fisher Scientific, Waltham, USA) and total RNAs were obtained according to the manufacturer’s protocol and then quantified and synthesized into cDNA using a iScript cDNA Synthesis Kit (Bio-Rad, Hercules, USA) for SALL4 expression or using a High-capacity cDNA Reverse Transcription Kit (Thermo Fisher Scientific, Waltham, USA) for miRNA expression. Real-time PCR was performed using SYBR Green Super Mixes (Bio-Rad, Hercules, USA). GAPDH and U6 were used as endogenous controls for normalization. MiRNA-specific primers were purchased from Ribobio (Guangzhou, China). Relative levels of expression were normalized and calculated using 2−ΔΔCt method. Information on the primers are listed in supplemental document 2.

### SALL4 expression in patient-derived samples

Patient samples were obtained with written informed consent in accordance with the Declaration of Helsinki and approval of the human research ethics committee of the South Eastern Sydney Local Health District (HREC ref# 17/295). Patient characteristics are available in Table 1. Bone marrow samples were collected prior to cycle 1 day 1 (C1D1) and following cycle 7 day 1 (C7D1) 6 cycles of subcutaneous administration of 5-azacytidine (Vidaza, Celgene), and mononuclear cells isolated using LymphoPrep (Axis-Shield). RNA was isolated with All-in-One DNA/RNA Mini-Preps Kit according to the manufacturer’s protocol (Bio Basic) and reverse transcribed using either a QuantiTect Revese Transcription Kit (Qiagen) or Maxima H Minus Reverse Transcriptase (Thermo Scientific) according to the manufacturer’s protocol. Quantitative PCR for SALL4 and GAPDH was performed using PowerUp SYBR Green Master Mix (Applied Biosystems) and a MX3000P thermocycler (Stratagene), and relative levels of expression calculated using the 2-ΔΔCt method. Information on the primers are listed in supplemental document 2.

**Table 1.**
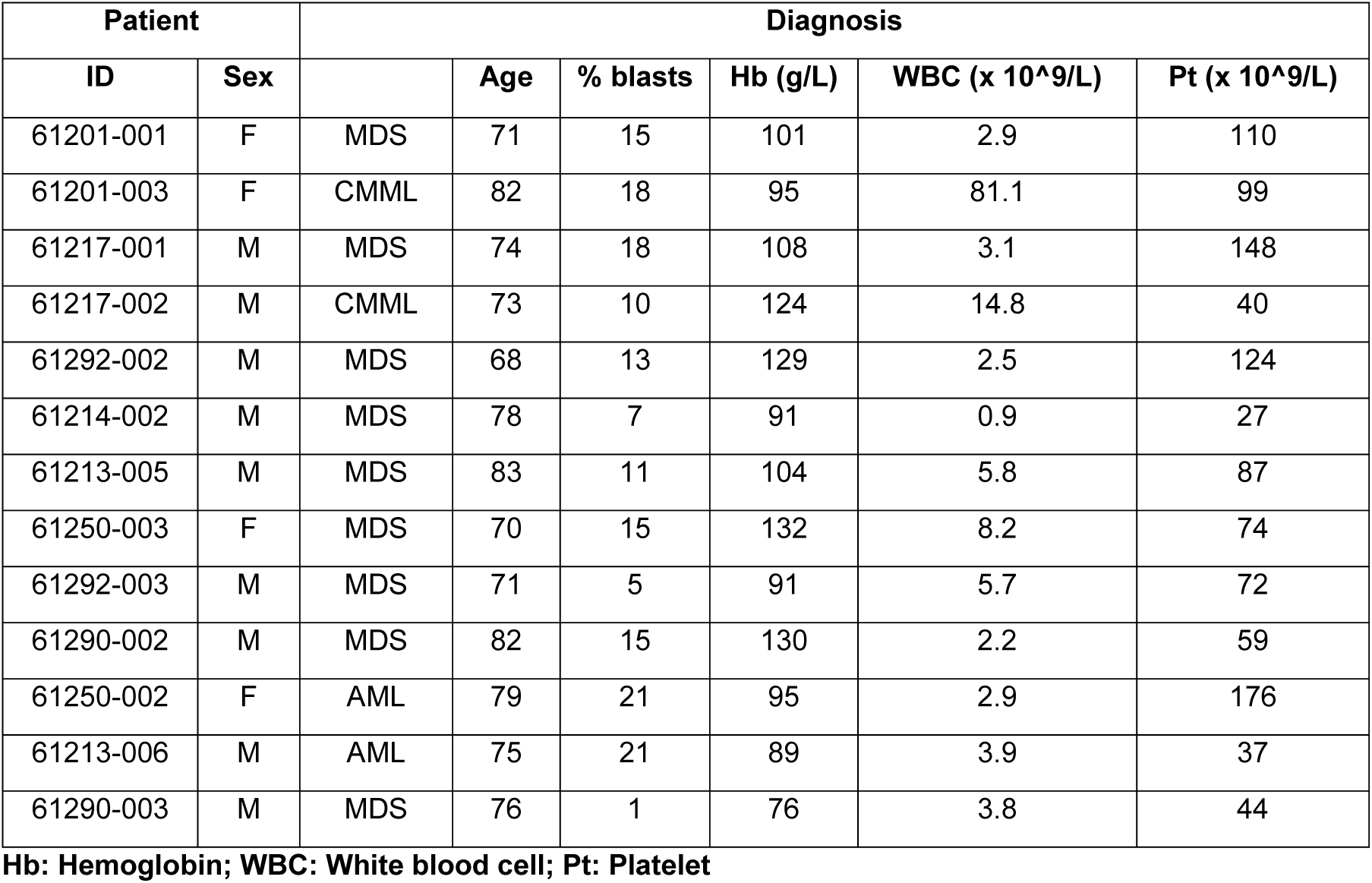
Patients’ characteristics.

| Patient |  | Diagnosis |  |  |  |  |  |
| --- | --- | --- | --- | --- | --- | --- | --- |
| ID | Sex |  | Age | % blasts | Hb (g/L) | WBC (x 10<sup>9</sup>/L) | Pt (x 10<sup>9</sup>/L) |
| 61201-001 | F | MDS | 71 | 15 | 101 | 2.9 | 110 |
| 61201-003 | F | CMML | 82 | 18 | 95 | 81.1 | 99 |
| 61217-001 | M | MDS | 74 | 18 | 108 | 3.1 | 148 |
| 61217-002 | M | CMML | 73 | 10 | 124 | 14.8 | 40 |
| 61292-002 | M | MDS | 68 | 13 | 129 | 2.5 | 124 |
| 61214-002 | M | MDS | 78 | 7 | 91 | 0.9 | 27 |
| 61213-005 | M | MDS | 83 | 11 | 104 | 5.8 | 87 |
| 61250-003 | F | MDS | 70 | 15 | 132 | 8.2 | 74 |
| 61292-003 | M | MDS | 71 | 5 | 91 | 5.7 | 72 |
| 61290-002 | M | MDS | 82 | 15 | 130 | 2.2 | 59 |
| 61250-002 | F | AML | 79 | 21 | 95 | 2.9 | 176 |
| 61213-006 | M | AML | 75 | 21 | 89 | 3.9 | 37 |
| 61290-003 | M | MDS | 76 | 1 | 76 | 3.8 | 44 |

**Hb: Hemoglobin; WBC: White blood cell; Pt: Platelet**

### Western blot analysis

Cells were washed by cold PBS and treated with lysis buffer (150 mM NaCl, 50 mM Tris pH7.5, 1 mM EDTA, 0.5% Triton X-100, 0.1% deoxycholic acid sodium salt) on ice for 30 mins. Then cells were scraped and after centrifugation lysate supernatant was collected and stored at −80°C. The BCA Assay kit (Bio-Rad, USA) was used to test protein concentration. Protein samples were denatured and then separated on SDS-PAGE and transferred to PVDF membrane (Millipore, Burlington, USA). After blocking by non-fat milk for 1 hour, membranes were incubated overnight at 4°C with primary SALL4 antibody from Santa Cruz Biotechnology (Cat. EE-30, Dallas, USA), PARP antibody from Cell Signaling Technology (Cat. 9542S, MA, USA) and β-actin antibody from Sigma-Aldrich (Cat. A1978, St. Louis, USA) with 1:1000 dilution. Then after washing 3 times, membranes were incubated with anti-mouse HRP-conjugated secondary antibody from GE Healthcare (Cat. NA9311ML, Chicago, USA). The bands were analyzed by an Immobilon Western Chemiluminescent HRP Substrate (Millipore, Burlington, USA).

### MiRNA sequencing

Total RNA was extracted from the H661 cells after DMSO or ENT treatment (duplicate samples for each treatment) for 8 hours using TRIzol (Invitrogen, Carlsbad, USA) and a miRNeasy mini kit (Qiagen, Hilden, Germany), following the manufacturer’s instructions. The RNA samples were sent to the Molecular Biology Core Facilities of Dana-Farber Cancer Institute for library preparation and Illumina MINIseq Next Generation Sequencing. Raw data could be accessed at GSE153589. miRNAs expression with at least 2-fold change was considered as significantly changed after ENT treatment.

### ChIP-qPCR

Briefly, 10 million cells after indicated treatment were crosslinked with 1% formaldehyde, then lysed in buffer containing 1% SDS, and spun at 20,000RPM to isolate the chromatin. Sonication was performed in lysis buffer containing 0.1% SDS. Protein A/G PLUS-Agarose (Santa Cruz Biotechnology, Dallas, USA) were conjugated with H3K27ac or IgG antibody (Cell Signaling Technology, Danvers, USA) and incubated with sonicated chromatin overnight at 4 degrees. Washes (0.1% SDS lysis buffer, LiCl wash buffer, and tris-EDTA) were performed and the chromatin-protein complex was reverse crosslinked by heating at 65 degrees with proteinase K (8ug/mL). The chromatin was incubated with RNaseA (8ug/mL) before it was extracted, purified with phenol chloroform (pH 8), and precipitated by ethanol. qPCR was performed using SYBR Green Super Mixes (Bio-Rad, Hercules, USA). Information on the primers are listed in supplemental document 2.

### Cell immunofluorescence staining

H661 cells were washed twice with cold PBS and then fixed in 4% paraformaldehyde/PBS for 10 min. After washing with PBS three times, cells were incubated with 0.2% Triton X-100 for 10 min. Then the cells were incubated with primary antibody against SALL4 at 4°C overnight. Antibody was applied in PBS containing 1% bovine serum albumin, followed by incubation with secondary antibodies for 2 hours at 37°C. Images were obtained using a laser microscope (Olympus, Japan).

### Cell viability and apoptosis assay

The Cell Counting Kit-8 (CCK8, Dojindo, Japan) was used to detect cell viability. Briefly, 1×10^3^ cells after the indicated treatment were seeded into 96-well plates and cultured for 5 days. Then 10 μL of CCK-8 solution was added to each well. After 3 hours culturing, the absorbance at 450 nm was measured using a spectrophotometer. To determine cell apoptosis, the Apoptosis Detection Kit was used (BD Pharmingen, Bedford, USA), and cells were washed and resuspended in binding buffer, followed by staining with Annexin V and propidium iodide for 30 min prior of flow cytometry analysis.

### Dual-luciferase reporter assay

The wild-type 3’UTR region of SALL4 mRNA or a mutant without the miR-205 binding site (Figure 4E) was amplified using PCR and cloned into the pGL3 vector (Promega, Madison, USA). HEK 293T cells were seeded into 24-well plates, then co-transfected with the indicated vectors and miR-205 mimics or the miR-negative control using Lipofectamine 2000 according to the manufacturer’s protocol. Finally, luciferase activities were measured using the dual-luciferase reporter gene assay kit (Promega, Madison, USA).

### Bisulfite treatment and sequencing

SALL4 Exon 1 region methylation status was assessed using bisulfite sequencing or pyrosequencing. In brief, 1 μg of genomic DNA extracted using the PureLink Genomic DNA Mini Kit (Invitrogen) was bisulfite-converted by using the Epimark Methylation kit (NEB). Each experiment included non-CpG cytosines as internal controls to detect incomplete bisulfite conversion of the input DNA. For manual methylation tests on H1299 cells, PCR products were gel-purified (Qiagen) from the 1.5% TAE gel and cloned into the pMD20T vector (Takara Bio) for transformation. The cloned vectors were transformed into ECOS 101 DH5α cells and miniprep was performed to extract plasmids. Sequencing results were analyzed using BiQ analyser software. The number of clones for each sequenced condition was 10.

Pyrosequencing of the SALL4 gene in H661 cells was performed by EpigenDx, Inc. (Hopkinton, MA, United States). The PCR product was bound to Streptavidin Sepharose HP (GE Healthcare Life Sciences), after which the immobilized PCR products were purified, washed, denatured with a 0.2 μM NaOH solution, and rewashed using the Pyrosequencing Vacuum Prep Tool (Pyrosequencing, Qiagen), as per the manufacturer’s protocol. Pyrosequencing of the PCR products was performed using 0.5 µM of sequencing primer on the PSQ96 HS System (Pyrosequencing, Qiagen) according to the manufacturer’s instructions. The mean methylation level was calculated using methylation levels of all measured CpG sites within the targeted region of SALL4 gene.

### Tumor cell implantation and treatment

All experimental procedures involving animals were conducted in accordance with the institutional guidelines set forth by the Children’s Hospital Boston (CHB animal protocol number 11-09-2022). Eight to ten-week-old non-obese diabetic scid gamma (NSG) mice were housed in a specific pathogen-free facility. One million H1299 cells subjected to different drug treatments were injected subcutaneously into the hidnlimbs on both sides. Cells were resuspended in 0.1 mL of PBS and then mixed with 0.1 mL of Matrigel. Tumors were harvested and weighed after 30 days. For drug treatment, after 10 days of H1299 cells implantation, mice received an intraperitoneal injection of either vehicle or DAC (2.5 mg/kg) for 5 days. Then an intraperitoneal injection of either vehicle or ENT (2.5 mg/kg) were given for the following two weeks (5 consecutive injections per week). Tumor volume was calculated by using the formula: Tumor volume=length x width^2^ / 2.

### Statistical analysis

Statistical analysis was performed by GraphPad Prism 8 (GraphPad Software, San Diego, USA). All experiments were conducted for triplicate. All data were presented as mean ± SD. The data were analyzed by Student t test. P < 0.05 was considered as statistically significant.

