## Supplement data for "Induction of a SALL4-dependency for targeted cancer therapy"

**Supplement Figure 1. SALL4-dependency could be established in its negative cells.** (A) Cancer cell lines with high SALL4 expression (top 100) were more dependent on SALL4 compared to those with low SALL4 expression (top 100). A score of 0 in Y-axis is equivalent to a gene that is not essential whereas a score of -1 corresponds to the median of all common essential genes. Data generated from depmap.org/portal/(*1*). (B) Cell viabilities of SALL4 positive SNU398 after knocking down SALL4 by siRNA. (C) Cell viabilities of GFP or SALL4-SNU387 cells after indicated knocking down treatment. (D) Hierarchical clustering and (E) similarity matrix of genes significantly changed after SALL4 overexpression in SNU387 cells (over 2-fold) based on Pearson correlation, data generated from GSE114808. (F) GO analysis of significantly changed genes upon SALL4 overexpression in SNU387 cells (over 2-fold). N=3.

**Supplement Figure 2. DAC treatment did not affect SALL4 expression in SALL4 positive cells.** (A) mRNA and (B) protein levels of SALL4 in H661 cells after 5 days treatment of DAC or DMSO. n.s. means P>0.05.

**Supplement Figure 3. ENT treatment decreased SALL4 expression.** (A) mRNA level of SALL4 in H661 cells after indicated treatment. (B) Protein and (C) mRNA levels of SALL4 in SNU398 cells after indicated ENT or DMSO treatment. (D) Pyrosequencing results of DNA CpG island methylation levels within SALL4 first exon after DMSO or ENT treatment. (E) ChIP-qPCR of SALL4 promoter region and non-specific control region after ENT treatment, IgG was used as normalized control. (F) The plot of SALL4 mRNA levels in different lung cancer cell lines and their corresponding IC50s against ENT, SALL4 level in H1299 cells was normalized as 1. n.s. means P>0.05, *P<0.05, **P<0.01, ***P<0.001, N=3.

**Supplement Figure 4. ENT inhibited SALL4 expression via miR-205.** (A) miR-205 predicted targets enrichment in KEGG pathway. (B) Represent images of SALL4 immunofluorescent staining on H661 cells after overexpression of scramble or miR-205 by GFP expressed plasmids. Arrows indicated cells transfected with scramble control (white) or miR-205 (yellow) overexpression plasmid. (C) mRNA and (D) protein levels of SALL4 in ENT-treated H661 cells after overexpression of miR-205 inhibitor or non-specific control. (E) Cell viability of ENT-treated H661 cells after indicated treatment for 5 days. NC was used as normalized control. (F) Flow cytometry and (G) statistic results of H661 cells after indicated treatment Annexin V and PI double staining, Apoptotic/Dead cell is positive for both or either, Live cell is negative for both. *P<0.05, **P<0.01, N=3.

**Supplement Figure 5. Sequential combination treatment in SALL4 negative cells.** (A) Relative level of miR-205 in DMSO or DAC-treated H1299 cells after indicated treatment. (B) IC50 of ENT for SNU387 cells after 5 days treatment of indicated DAC or DMSO. (C) and (D) Cell proliferation of K562 and HL60 cells with indicated treatment. *P<0.05, **P<0.01, ***P<0.001, N=3.

**Supplement Figure 1**

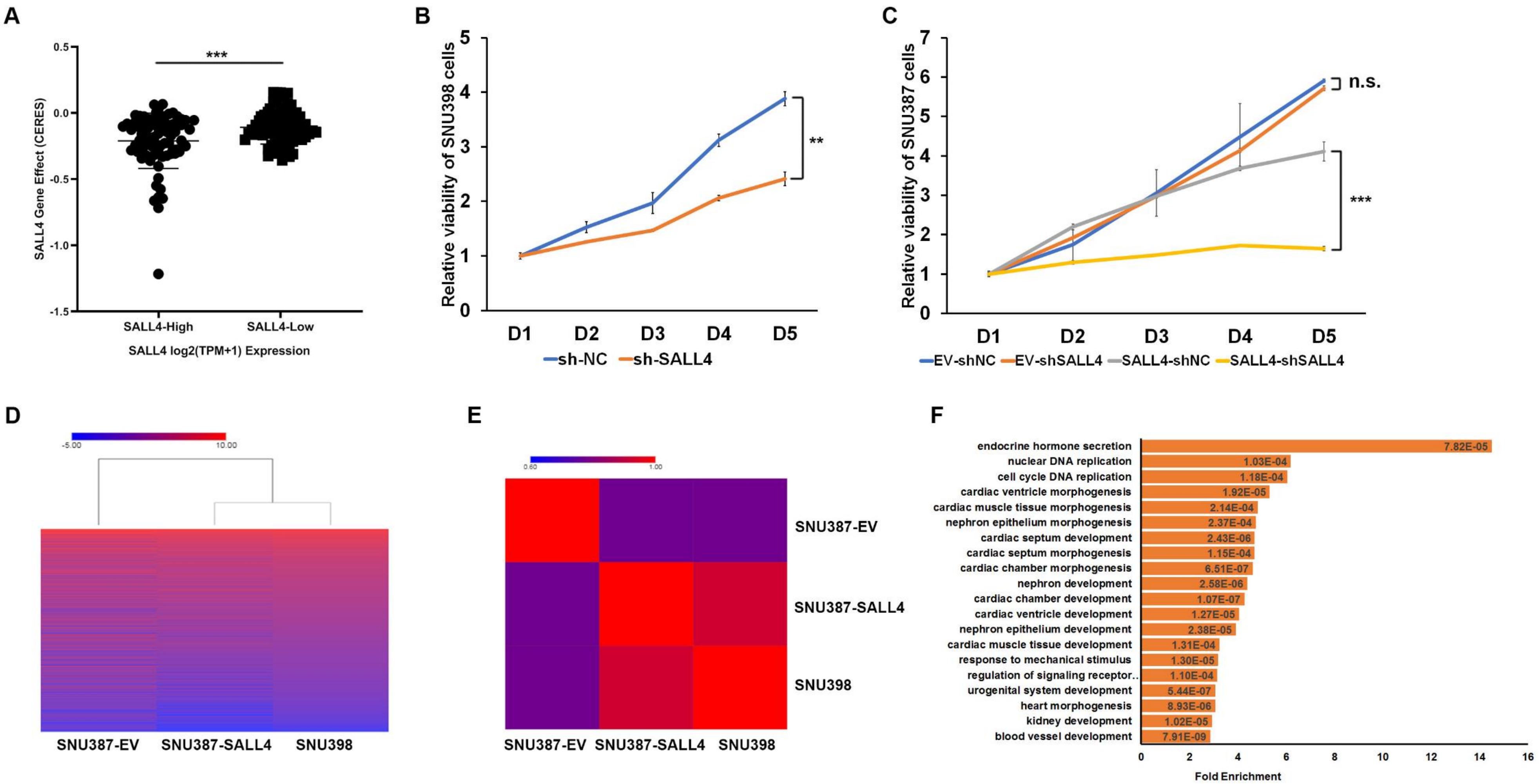

**Supplement Figure 2**

A

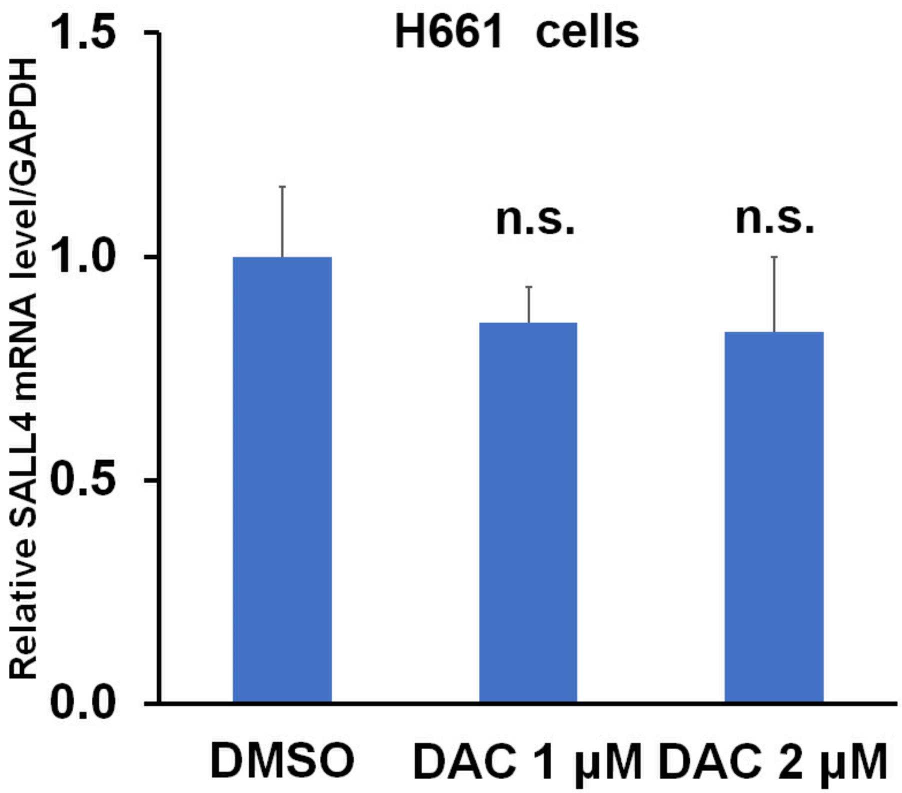


B

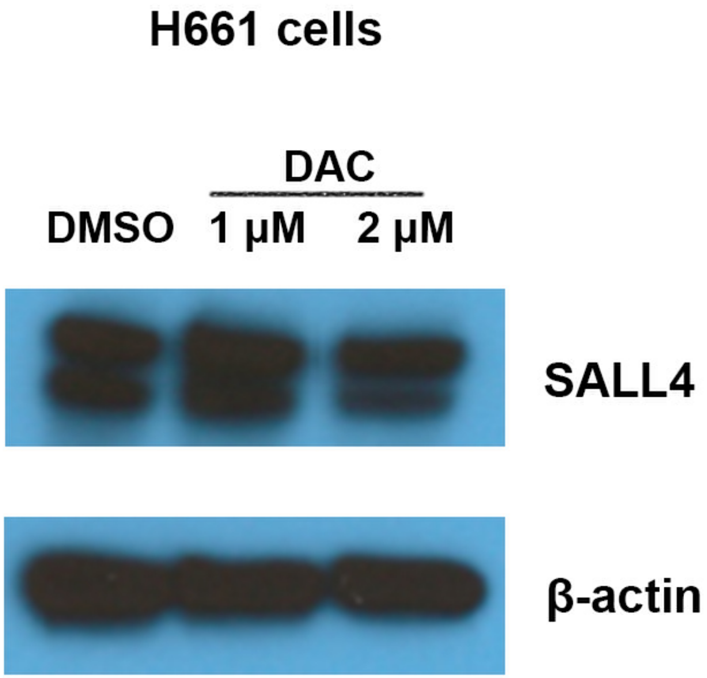

**Supplement Figure 3**

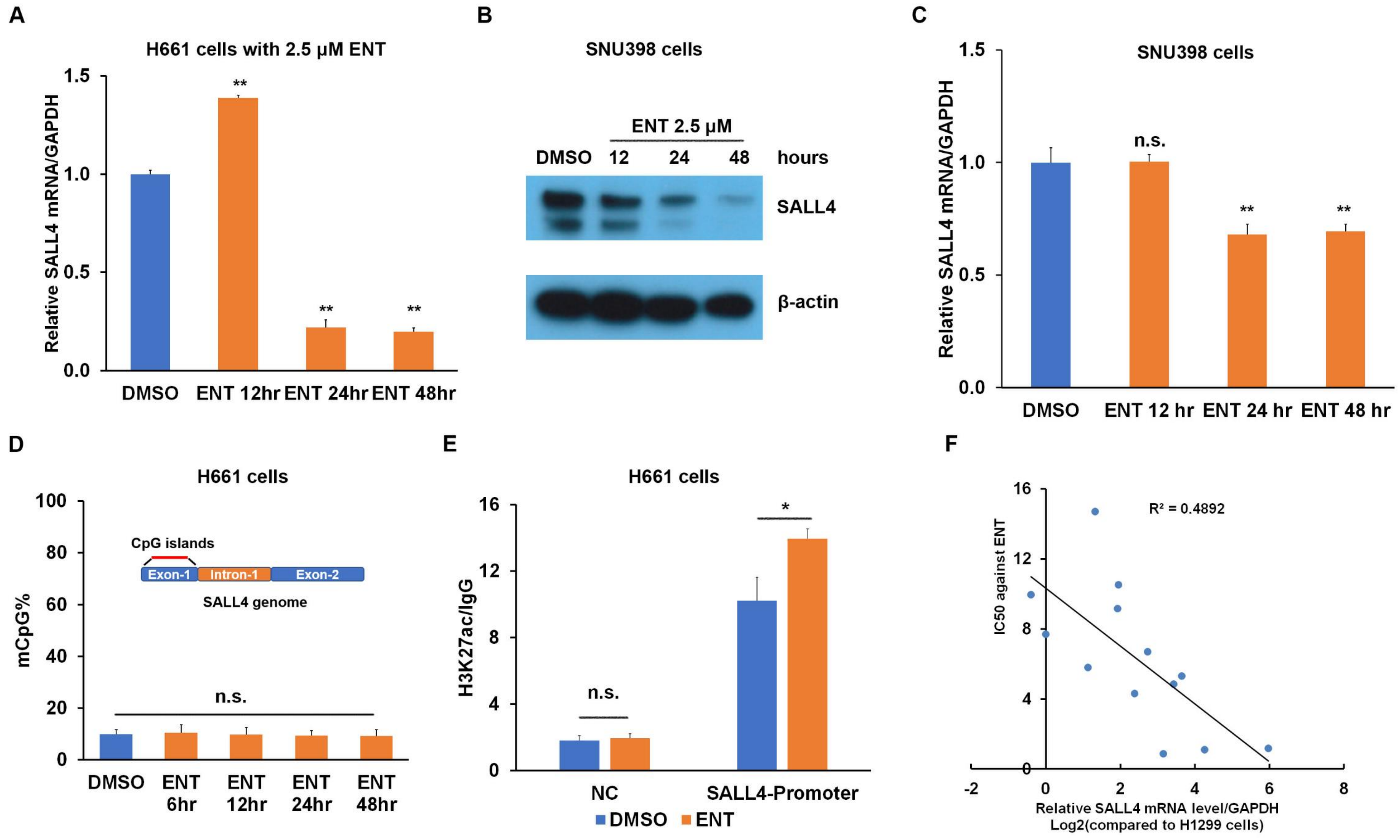

**Supplement Figure 4**

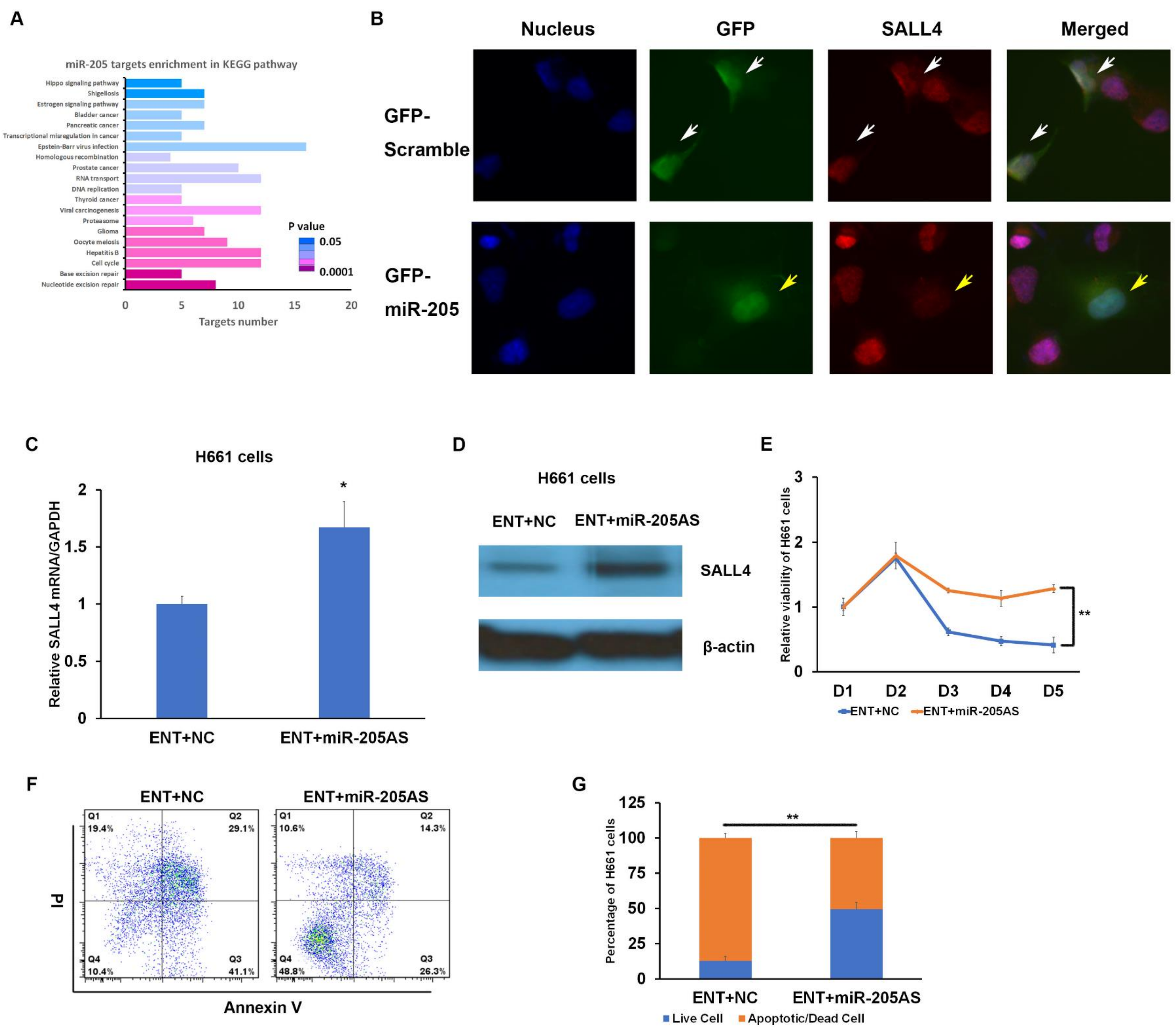

**Supplement Figure 5**

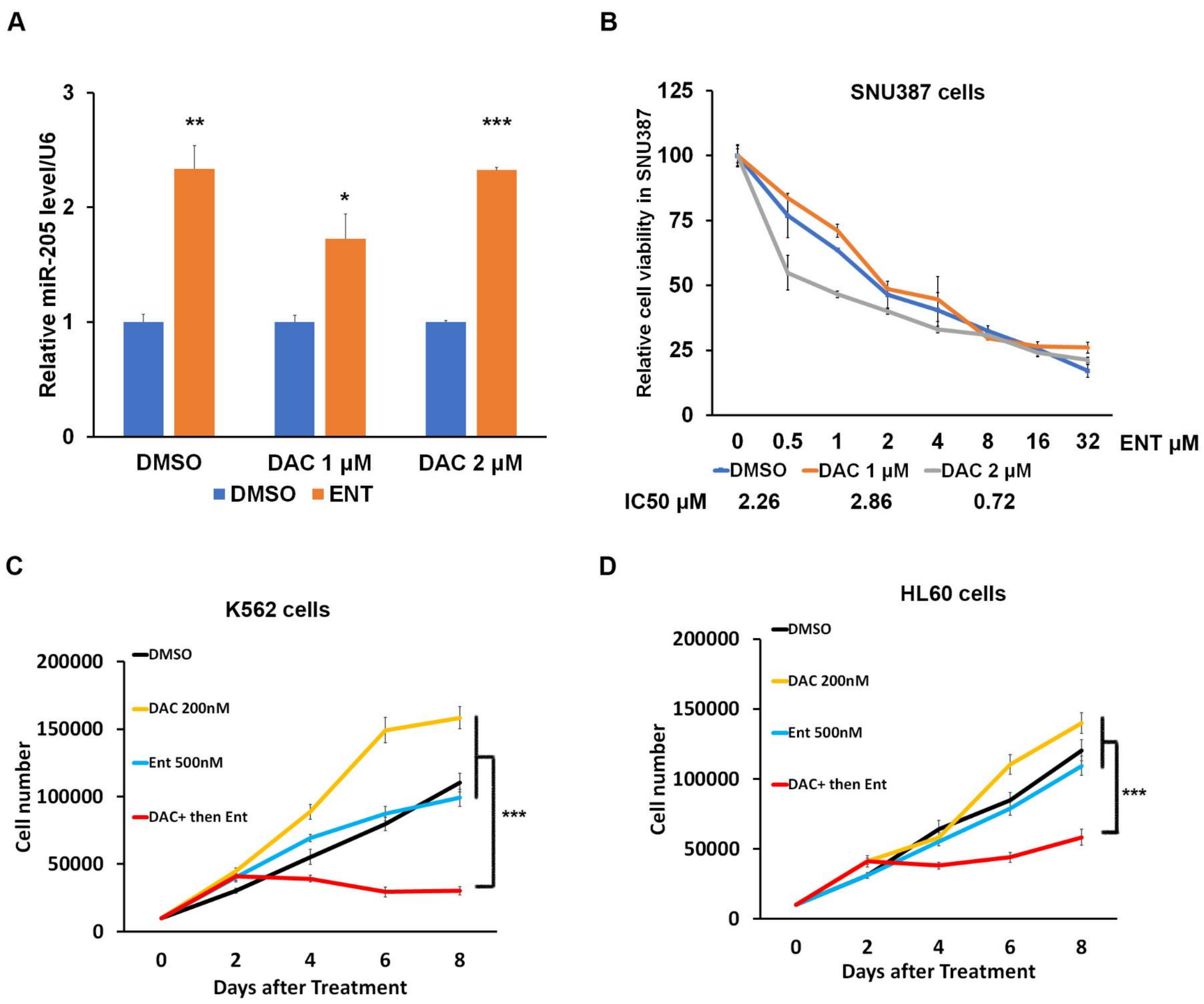

**Supplement document 1. List of primers used in the experiments.**

**Supplement document 2. List of significantly changed miRNAs after ENT treatment in H661 cells.**
